# Temperature accounts for the biodiversity of a hyperdiverse group of insects in urban Los Angeles

**DOI:** 10.1101/568212

**Authors:** Terrence P. McGlynn, Emily K. Meineke, Christie A. Bahlai, Enjie Li, Emily A. Hartop, Benjamin J. Adams, Brian V. Brown

**Author notes:** Authors contributed equally to this work (Terrence P. McGlynn, Emily K. Meineke).

## Abstract

The urban heat island effect is a worldwide phenomenon that has been linked to species’ distributions and abundances in cities. However, effects of urban heat on biotic communities are nearly impossible to disentangle from effects of land cover in most cases because hotter urban sites also have less vegetation and more impervious surfaces than cooler sites within cities. We sampled phorid flies, one of the largest, most biologically diverse families of true flies (Insecta: Diptera: Phoridae), at 30 sites distributed within the central Los Angeles Basin, where we found that temperature and the density of urban land cover are decoupled. Abundance, richness, and community composition of phorids inside urban Los Angeles were most parsimoniously accounted for by mean air temperature in the week preceding sampling. Sites with intermediate mean temperatures had more phorid fly individuals and higher richness. Communities were more even at urban sites with lower minimum temperatures and sites located further away from natural areas, suggesting that communities separated from natural source populations may be more homogenized. Species composition was best explained by minimum temperature. Inasmuch as warmer areas within cities can predict future effects of climate change, phorid fly communities are likely to shift non-linearly under future climates in more natural areas. Exhaustive surveys of biotic communities within cities, such as the one we describe here, can provide baselines for determining the effects of urban and global climate warming as they intensify.

## Introduction

Urban development is accelerating with uncertain effects on biodiversity. While many species do not persist in urban areas, cities can support a surprising range of native and even threatened taxa [1,2]. Thus, determining conditions within cities that affect species persistence is increasingly a focus of ecological research from fundamental and conservation perspectives [3–5]. However, isolating specific drivers of biodiversity in cities has proven difficult because organisms in cities experience a range of novel conditions that may alter their abundances and distributions [6–8]. Therefore, for most animal taxa, specific mechanisms driving community assembly in cities remain unknown.

The urban heat island effect is a prevalent phenomenon in cities, and growing evidence shows that urban heat can alter species richness, abundance, and community composition [9–15]. Urbanization causes cities to be as much as 12°C hotter than adjacent areas [16], which is on par or above warming anticipated by the Intergovernmental Panel on Climate Change over the next several decades [17]. In certain cities, urban warming can also operate at local scales, creating thermal mosaics within the urban matrix (e.g., [18–21]). Despite the short history of research on the biotic effects of urban heat, researchers have found important patterns across diverse taxa [7,10–13,22–24]. For example, remnant native plant communities in urban environments may be altered under warming conditions, favoring more xerophilic species [25,26].

Because temperatures in cities match or exceed those expected under future climate change, researchers have suggested that thermal gradients within cities might allow us to predict biotic responses to the future climate warming [27,28]. Cities might be useful proxies for climate warming because urban heat has been in place for decades to centuries, and large scale, controlled warming experiments in more natural areas are often impractical (but see [29,30]). However, urban heat might not be an appropriate proxy for broader climate warming because other aspects of urbanization might also have strong effects on species. Perhaps most importantly, land cover (impervious surfaces, vegetation) and urban heat tend to covary, making it impossible to separate their effects on biological processes. Hot urban environments often have more impervious surface, less vegetation cover, and lower vegetation complexity [31–33]. While researchers have used various useful approaches to determine effects of urban warming alone – e.g., laboratory chamber experiments [11,18] – actually separating effects of land cover and temperature in the city could provide insight into whether biotic responses are more attributable to temperature or other aspects of urbanization. In coastal cities, urban temperatures are often decoupled from landcover, such that sites that are highly urbanized are not necessarily hotter than surrounding sites that are less urbanized because of winds entering from the coast [40]. This offers an experimental opportunity to separate the ecological effects of urbanization and temperature.

Insects are highly responsive to temperature, are a foundational component of terrestrial biodiversity, and provide a range of services and disservices within cities [34,35]. As insects are ectotherms, they have elevated metabolic and reproductive rates in response to warming until their thermal maxima are reached [14]. One of the most abundant animals in terrestrial environments are phorid flies [36,37], which are responsive to thermal conditions, but also feed on a wide range of resources and develop and occupy a tremendous variety of microhabitats [38]. Cities can support hyperdiverse communities of phorid flies, with dozens of species recently described from central Los Angeles [39,40]. With a small body size (0.4-6 mm) and presumably short dispersal distances, we would expect phorid fly biodiversity to finely track microclimatic conditions in the urban environment, relative to less ephemeral or larger-sized organisms.

Here we evaluate the spatial and temporal predictors of phorid fly biodiversity within urban Los Angeles, CA, USA, hereafter we refer to as L.A. In L.A., urban temperatures are decoupled from land cover, allowing us to investigate the effects of impervious surface,vegetation cover, and temperature,in a system where these aspects of the urban environment are not highly correlated [41]. In this project, species were sampled exhaustively [42], and 30 new species of flies were described from L.A. from this dataset in 2015 [39]. We leverage the complete documentation of this diverse group to determine effects of urban land development and climate in a city where we found these variables are uniquely decoupled. We sampled phorid flies and site environmental conditions in 30 locations throughout a calendar year to evaluate biodiversity responses to thermal and urbanization gradients within the L.A. metropolitan area. By measuring temperature and moisture variables at a very fine scale to match the habitat occupied by the organisms [43], we achieve a biologically relevant understanding of how local climatic factors vary across an iconic urban habitat.

## Methods

### Study area

The Los Angeles metropolitan area is a highly urbanized region located at 34ºN along the west coast of North America, which has experienced rapid population growth and associated land development over the past 100 years. The climate and flora are characteristically “Mediterranean,” and biomes that have given way to development include coastal sage scrub, chaparral, and oak woodlands. Some habitats have only small fragments remaining, including coastal dunes and wetlands [44]. The climate of locations within the city can vary substantially from one another, because of differences in distance from the ocean, elevation, intensity of urbanization, and vegetation [41]. The heterogeneity of the landscape makes predicting climatic differences among sites in Los Angeles particularly difficult [45,46], which reinforces the need for site-specific weather records to reliably compare sites.

### Study design and insect sampling procedures

We placed a series of Malaise traps [47] (Townes lightweight model Sante Traps, Lexington, KY) in 30 sites throughout central L.A. (Fig. 1). The distribution of the sites was designed to capture a range of biotic and abiotic gradients in the urban environment as part of the BioSCAN (Biodiversity Science: City and Nature) project of the Natural History Museum of Los Angeles County (LACM). The initial findings from this sampling are described by Brown and Hartop [42], who provide a detailed description of each site featured in the study. In a survey, participants whose homes were included in the study were asked if they used pesticides in their yards in areas close to where the traps were located. The survey revealed that none of the sites were treated with pesticides regularly, and only a few hosts used small quantities of pesticides for local control on rare occasions, such as neem oil on a few plants. We decided this incidental treatment would not appreciably affect biodiversity within the yards included in this study.

**Figure 1.**
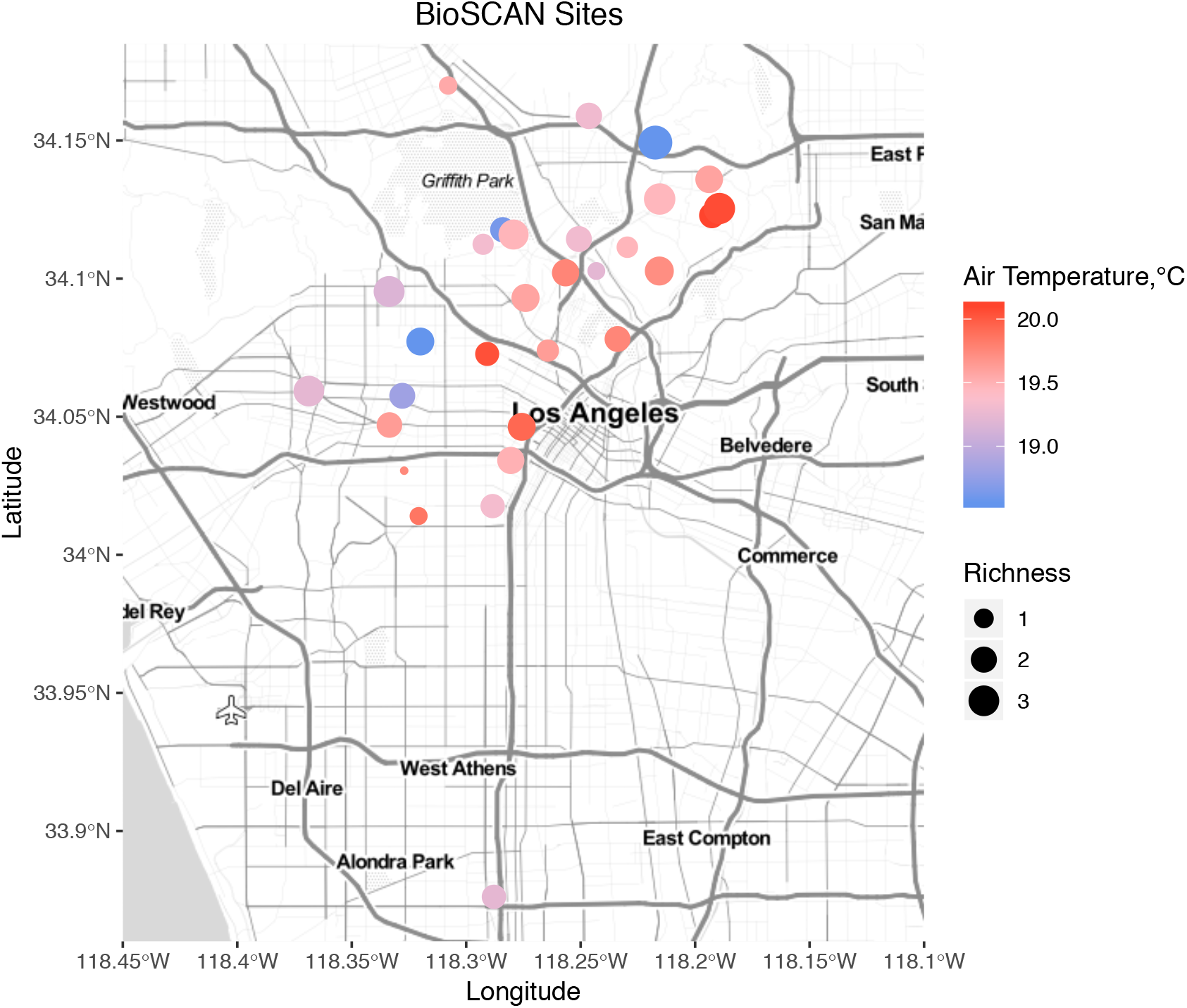
Map of BioSCAN sites where phorid flies and climatic variables were sampled. Dot size represents mean daily phorid fly species caught in traps, and color represents mean annual air temperatures.

For each of twelve sampling periods (approximately the first week of each month in 2014), we collected and identified all phorid flies in samples to species, resulting in a total of 42,480 specimens, . Vouchers are deposited in the LACM. Over this year of sampling, the fauna of 99 species was essentially sampled to completion, as richness estimators predicted that additional sampling would be expected to yield perhaps one additional new species [42]. We are confident that this sampling regime represents nearly all the species in this lineage and locality that would be captured using this sampling approach.

### Abiotic data collection and processing

We continuously recorded air temperature, soil temperature, and relative humidity at each site using a weather station adjacent to each trap (Onset HOBO U30 Station, Bourne, MA).

Additional details about abiotic data are in the electronic supplementary material.

### Statistical Analyses

#### Correlations between environmental and climatic predictors

To determine the relative contribution of urbanization and topography on microclimates across our study region, we used two simple linear models to test whether impervious surface and/or elevation were predictive of mean average annual air temperature at our sampling sites. In each model, mean average annual temperature was the response variable, and impervious surface or elevation was the sole predictor. We also evaluated whether differences in temperatures across sites were associated with a coastal effect from the Pacific Ocean. Our hypothesis was that urban sites further away from the coast would have warmer mean temperatures [50]. To test this, we also used a simple linear model, with distances from our sites to the Pacific shoreline as the predictor and mean average annual air temperature as the response variable.

#### Phorid fly abundance, richness, and evenness

We first calculated four response variables that were each used as the responses in the modeling framework described below. First, we calculated the total number of individuals caught per trap per day (abundance), species richness, and Pielou evenness. Because the traps were in place for slightly different amounts of time during some sampling periods, we divided each response variable by the number of days a trap was left out, i.e., the total amount of time flies had access to a trap. One species, *M. agarici*, constituted a substantial number of specimens in samples at many sites (and about one-quarter of all individuals collected). Therefore, we included total individuals of this species captured per trap per day as an additional response variable. (Many of the species in this study were only recently described and their biology remains poorly known, and the current state of knowledge [51] prevents us from using taxon-specific data, such as phylogeny, diet, as factors in the models described below.)

As a preliminary step, we used model selection to minimize overfitting in the final models. Specifically, we used model selection to identify the most parsimonious independent variables describing effects of temperature, humidity, and urbanization on each response variable. For each response, we built a series of linear mixed effects models in the *nlme* package in R [52]. In each case below, we compared models with tightly correlated predictors describing similar aspects of the urban environment and selected the parameter in the model with the lowest AICc score to include in full models used for inference, i.e., to choose the response variable most closely associated with the response. For each response variable, we compared three sets of models. One set included mean RH (relative humidity), maximum RH, minimum RH, and no humidity predictor. The second set of models selected from included mean temperature, maximum temperature, minimum temperature, and no temperature predictor, and the final set compared mean soil temperature, mean maximum soil temperature, mean minimum soil temperature, and no temperature predictor. All climatic predictors represented average conditions one week before sampling to represent the conditions most likely to affect phenology [53]. We decided, depending on the shape of the response, whether to include a squared term to account for non-linear responses of phorid flies to environmental variables. In all models built for final model selection, we included latitude, longitude, and distance to the nearest natural, protected area were included as fixed effects, and site was included as a random effect to account for repeated sampling of flies at each site. To account for the composition of the matrix surrounding each study site and to describe urbanization, we compared models that included impervious surface cover, NDVI – each measured at a 50-m buffer as described above – and neither of these (null model).

After selecting parameters for each response variable (species richness, *M. agarici* abundance, total abundance, and evenness), we built one full model for each, for a total of four models. In these models, we included each parameter chosen to represent urbanization, temperature, and humidity, along with latitude, longitude, and distance to the nearest natural area as fixed effects. Site was included as a random effect in all models. To determine if the effects of temperature depended on water availability, and vice versa, we included an interaction between the best temperature predictor and the best relative humidity predictor; these were subsequently removed from all models because they were not significant. We did not include any interaction effects that were not associated with explicit *a priori* hypotheses, because these interaction effects often can be explained more directly by main effects of environmental variables, and including these variables would be redundant and reduce the power of analyses of our main effects. To determine if site proximity rather than environmental conditions may account for responses we observed, we performed an analysis to test if spatial autocorrelation was observed among sample sites. We examined abundance of phorid flies by date, for every date where >5 sites reported data, using Moran's I, applied to an inverse distance matrix of site co-ordinates as the weighting factor [54].

#### Community composition

In addition to the univariate community responses, we also conducted a non-metric multi-dimensional scaling (NMDS) analysis to examine patterns in community composition, and fit environmental vectors to gain insights into drivers of these patterns [55]. For this analysis, we used all captures of phorid flies at a given site, across the whole sampling period. We culled all singletons (species represented by a single sample throughout the entire study), a standard approach because the incidence of a singleton is indistinguishable from a spurious occurrence [56]. The NMDS was conducted on the Bray-Curtis distance calculated from the untransformed matrix of taxon-by-site using abundance values. Environmental fit vectors were selected iteratively by comparing the fit statistics (Global R^2^ and p-value) within a group of related, and auto-correlated, parameters (e.g., minimum, maximum and mean air temperatures, etc.).

## Results

### Correlations between environmental variables and urbanization

Measured mean annual temperatures at our study sites were independent of impervious surface and elevation (Fig. 2; impervious surface: y= −129+9x, F_1,28_=1.12, *p*= 0.30; elevation: y= −45+4.5x, F_1,28_= 0.02, *p*= 0.90), suggesting that neither urban land cover nor elevation drove urban temperatures. Therefore, we conclude that, while L.A. may have a largescale urban heat island effect, other unknown factors drive temperatures at the local scale. As expected, NDVI tracked impervious surface, though these values were not closely related to the distance to natural areas (Fig. S1). Contrary to expectations, temperatures measured at weather stations in urban backyards were not significantly associated with their distances from the coast (y= 1.90+1.83e^−5^, F_1,28_= 0.48, *p*= 0.49).

**Figure 2.**
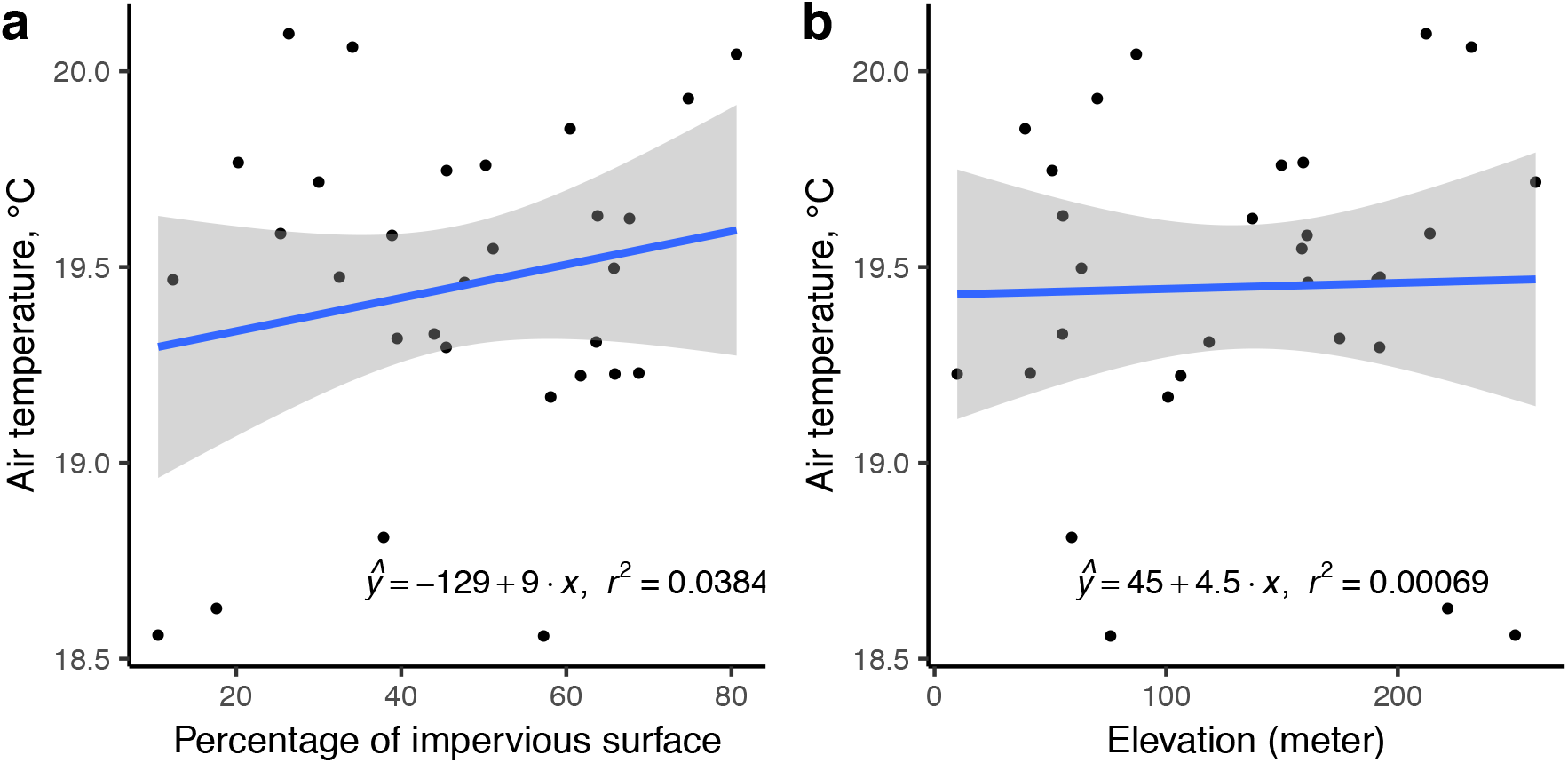
Thermal responses to landscape characteristics. Mean annual air temperature is not associated with a) impervious surface, nor with b) elevation.

### Mean and peak biotic responses

Abundance and richness of phorid fly communities throughout the city were best explained by air temperature (Table 1; Fig. 3). No other climatic parameters had significant predictive value, aside for a lesser effect of relative humidity on species richness (Table 1), and humidity slightly tracked temperature (Fig. S3). Phorid fly abundance and richness responses to environmental conditions were nonlinear, with peaks at intermediate mean weekly temperatures (Fig. S1). The factors affecting the abundance of the most common species, *M. agarici*, were the same factors affecting total abundance (Table 1). Evenness of phorid fly communities was weakly explained by mean minimum weekly temperature and distance to natural areas, such that phorid fly communities in areas with lower minimum temperatures and those that were further away from natural were more even (Table 1).

**Table 1.**
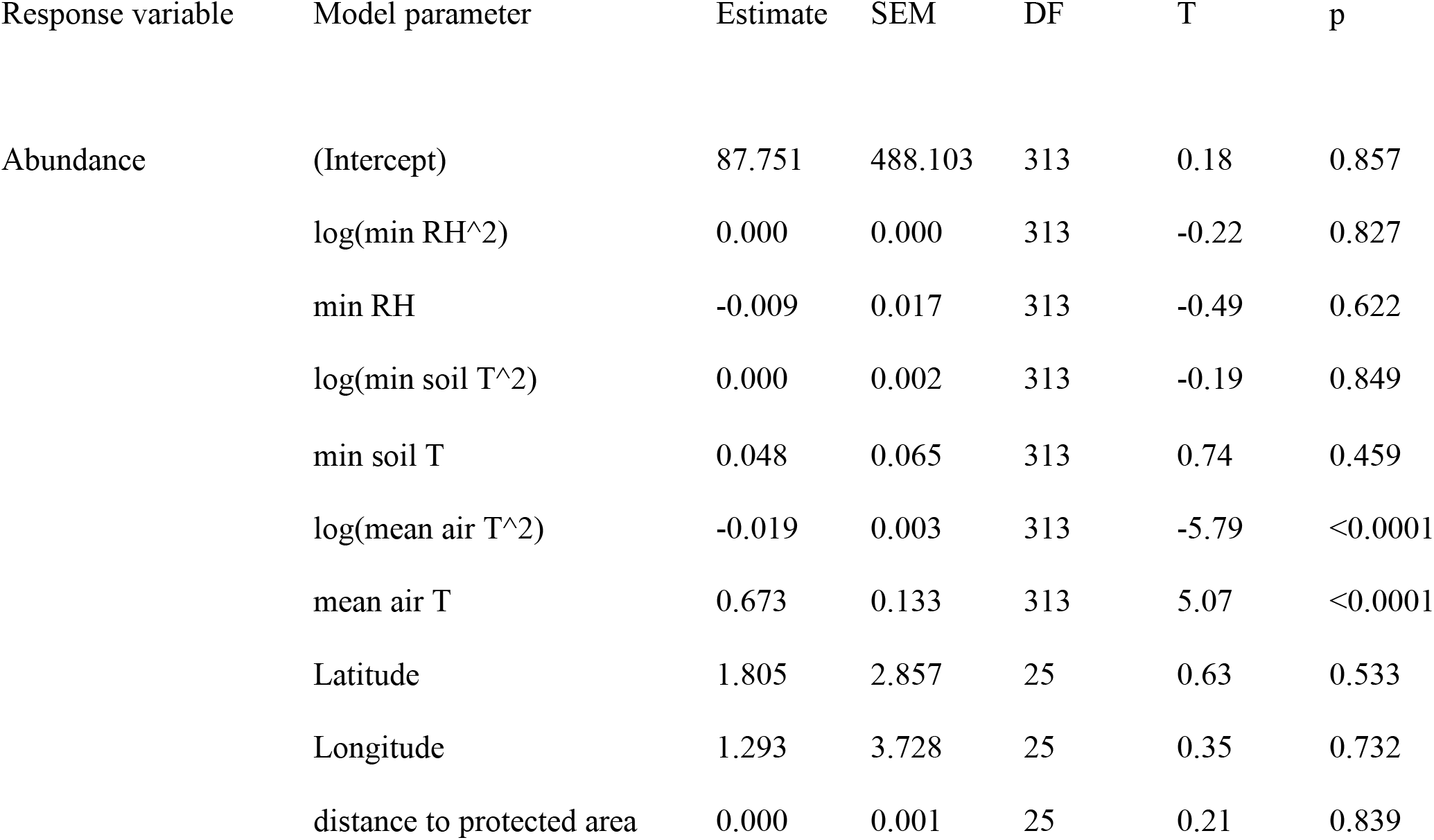
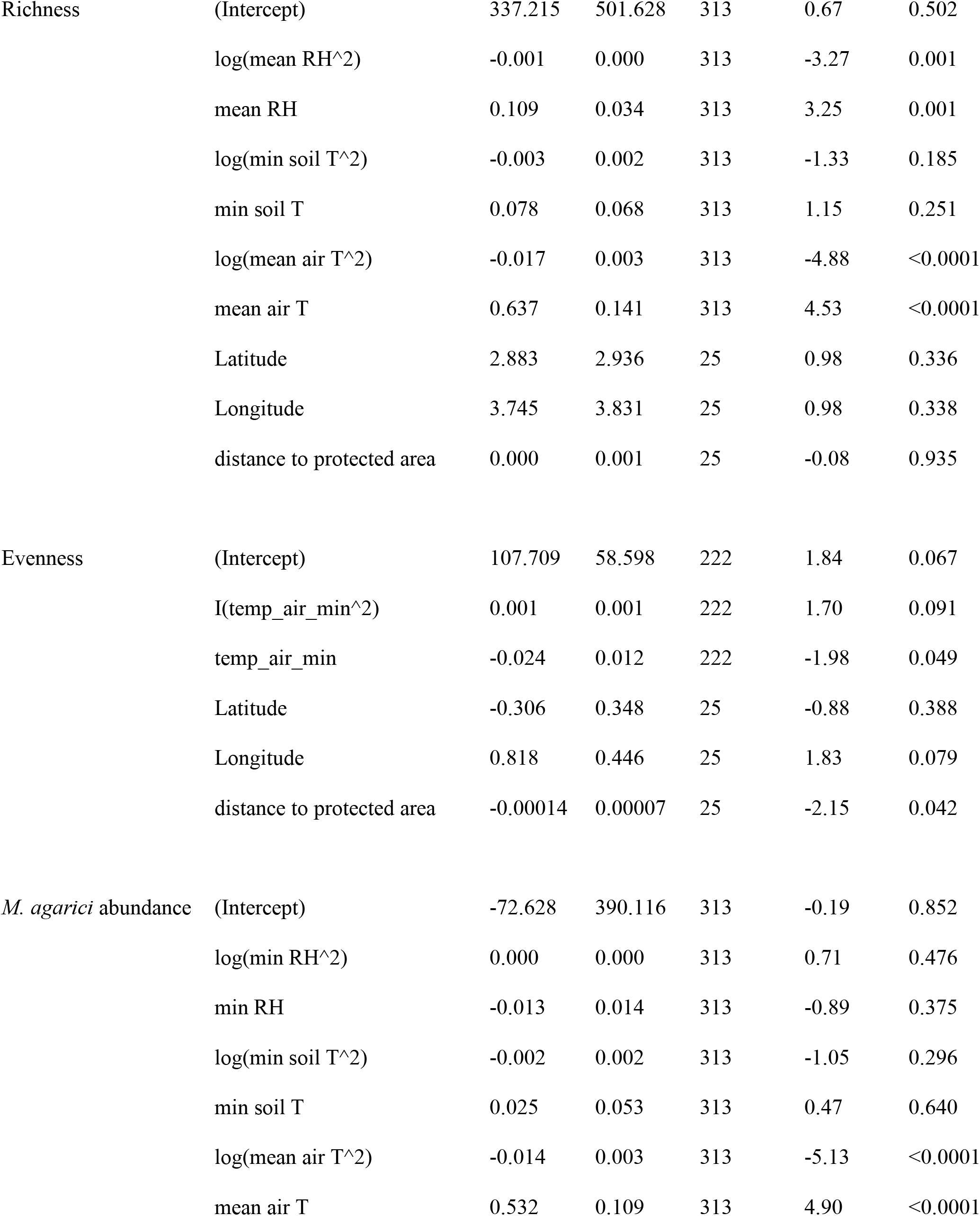
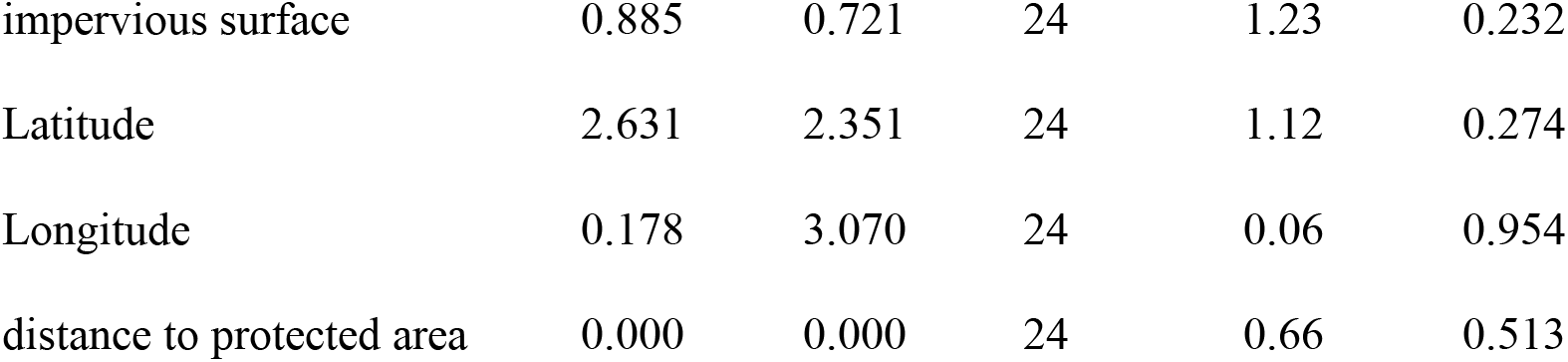
Full model evaluation of phorid fly biodiversity across 30 sites in urban Los Angeles.

**Figure 3.**
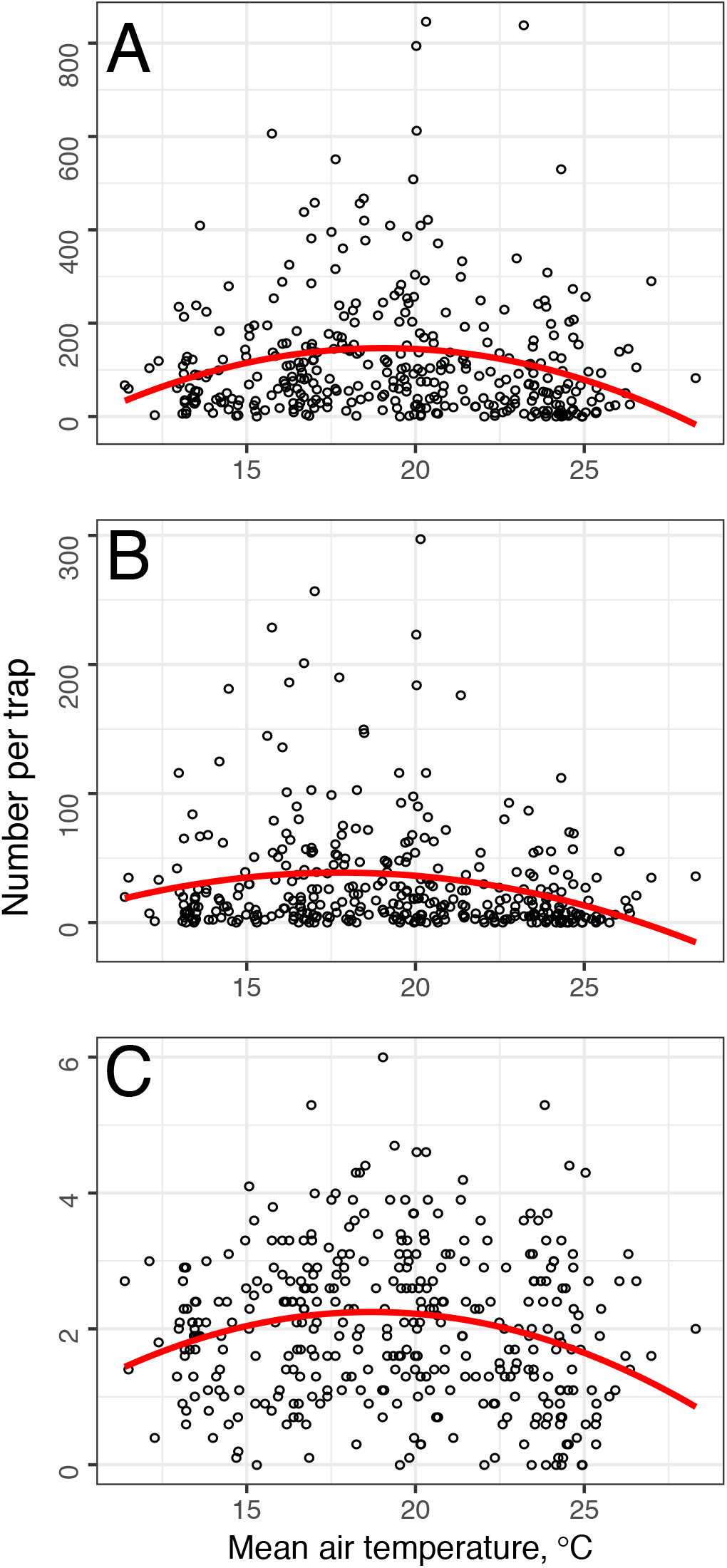
Phorid fly abundance and richness responses to temperature. a) Total abundance of phorid flies caught in each trap per sampling period, b) abundance of the most prevalent species, *M. agaraci*, and c) total species richness per sampling period. The x-axis represents mean air temperature the week prior to sampling, and regression lines represent best fits.

Latitude and longitude were associated with abiotic conditions (Figs. S4, S5), but spatial autocorrelation was limited. Among all dates where sufficient data existed for autocorrelation analysis (10 dates), one date (Week 6 of 2014) had significant spatial autocorrelation (*p*= 0.037), suggesting that autocorrelation is rare in this system and may have been observed by chance. Thus, after we accounted for spatial similarity of sites using latitude and longitude as described above, no additional correction for spatial autocorrelation was needed.

### Community composition

The effects of NDVI and mean minimum weekly temperature on species composition were orthogonal, with a much greater effect of temperature (Fig. 4). Mean minimum weekly temperature had the only significant vector, which also had the greatest magnitude (r^2^= 0.26, *p*= 0.02)

**Figure 4.**
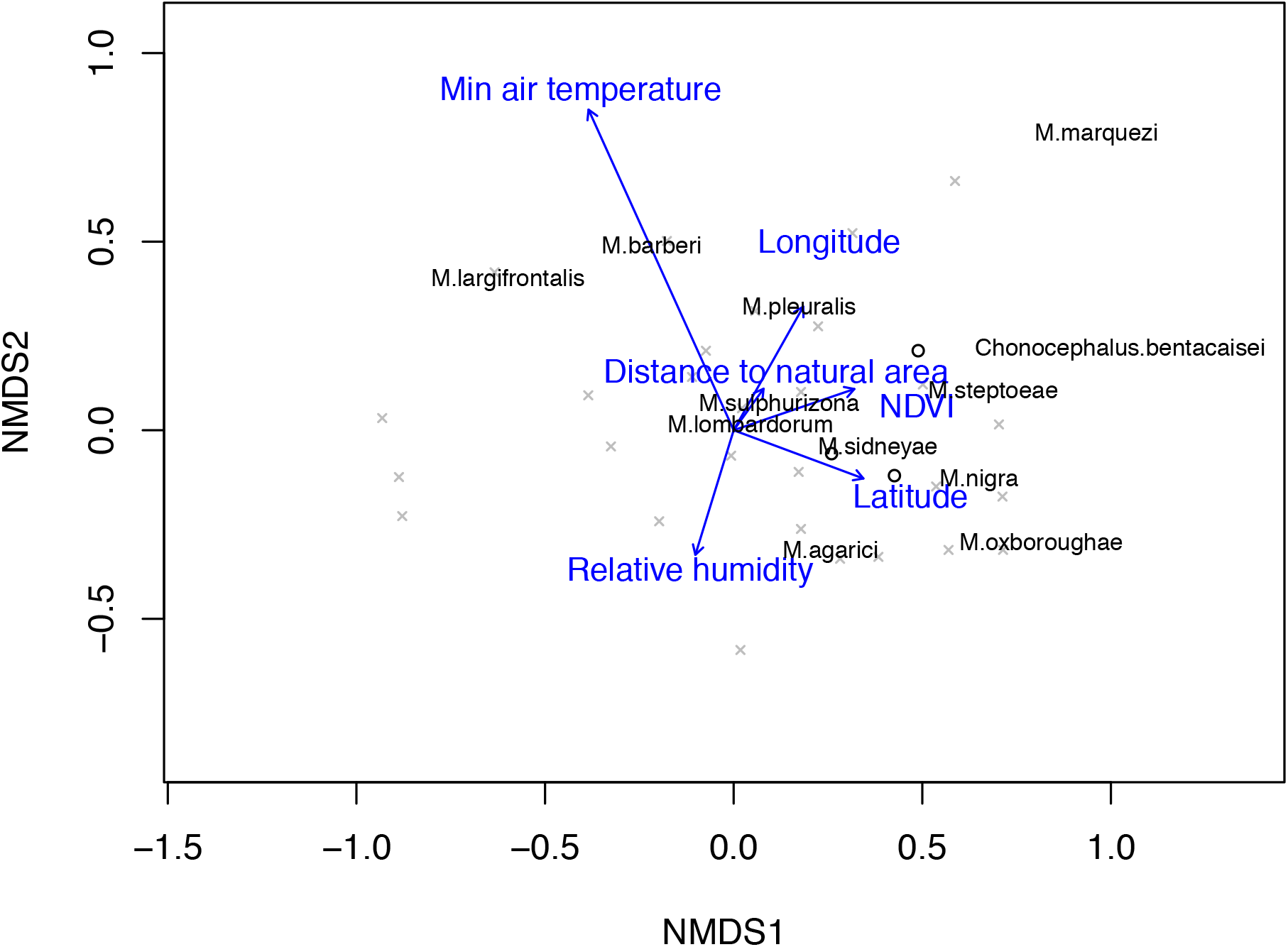
Non-metric multidimensional scaling of phorid fly communities from urban Los Angeles backyard sampling stations. The species are superimposed with environmental fit vectors for minimum air temperature (r^2^= 0.26, *p*= 0.02), normalized difference vegetation index within a 50-meter radius of the site (NDVI; r^2^= 0.03, *p*= 0.65), distance to the nearest natural area (r^2^= 0.01, *p*= 0.92), relative humidity (r^2^= 0.04, *p*= 0.63), latitude (r^2^=0.04, p=0.61) and longitude (r^2^= 0.04, *p*= 0.56). Most common species names are plotted (with captures >500 individuals). NMDS 2D stress= 0.13.

## Discussion

We found that urbanization and climate are uniquely decoupled across the L.A. Basin. Consistent with an earlier study [41], urban land cover does not influence local temperatures at the fine scale of our sampling Because of this decoupling, we were able to independently assess effects of local climate and urban land cover on phorid fly communities. We found that air temperature had the most robust influence on the assembly of the phorid fly community, but that different aspects of temperature were most closely associated with species abundance, richness, and evenness. Local impervious surface and vegetation cover (NDVI), which represent urbanization, did not outperform null models. We conclude that local climate, not urban land cover, is the strongest driver of phorid fly community assembly across Los Angeles.

Higher mean temperatures were associated with increased phorid fly richness, until around 20°C, where richness declined. We predicted that cooler sites would harbor more species because warmer areas would be associated with reduced persistence of heat-intolerant species. However, we find evidence that intermediate temperatures may support both heat-tolerant and - intolerant species, and thus most phorid species in L.A. Many of the species in our study (e.g., *M. halterata*, *M. nigra*, and *M. pleuralis*) are probably introduced from areas of northern Europe with cooler thermal conditions than L.A., which could also account in part for the loss of abundance at higher mean temperatures. However, in general, the relative contributions of non-native and native species to the patterns we observed are unclear. While we know that many species of phorids in L.A. are native based on their species interactions and/or distributions (i.e., found only in certain parts of North America in well-studied groups), knowledge about phorid fly distributions is inadequate to inform us to what extent non-native species contribute to the patterns we observe.

While species richness was tied to mean temperature, evenness was weakly explained by minimum temperature and distance to natural areas. The latter was included in the analysis because it is well established that some species have habitat requirements involving larger patches of land that are less urbanized. For example, the ant-decapitating guild of phorid flies are only found within and adjacent to natural areas because these are locations in L.A. where their hosts (species of *Camponotus*, *Crematogaster*, *Liometopum*, *Neivamyrmex*, *Pheidole*, and *Solenopsis*) are found. Elsewhere, the host ants are typically displaced by invasive Argentine ants [42,57]. We expected that species richness would drop with distance from natural areas, as certain native species would be removed from the species pool. Instead, we found that species evenness might be the result of a more complex process, in which communities become less even further from natural areas, but not because of the loss of species from communities – distance from natural areas did not predict richness – but rather because relative abundances may shift. Our results suggest that a subset of species may benefit from living further from natural areas, where perhaps there is less competition from species that may locally disperse into more urbanized environments from source populations in more natural areas. While we predict this may be the cause of less even communities further from natural, protected areas, further studies on population genomics and niche overlap of phorid flies are needed to determine mechanisms driving this pattern. In addition, sites with lower minimum temperatures support compositionally different and more even communities than areas with higher minimum temperatures. We may observe this pattern because a few species in the region have disproportionate fitness benefits from warmer minima.

Species richness was also higher at intermediate relative humidities, and we suspect this results from the benefits and drawbacks of wet climates for insects. Throughout the year, many species showed large spikes in abundance. Among those with known life histories (as listed in Fig. 4: *Chonocephalus bentacaisei*, *M. agarici*, *M. halterata*, *M. marquezi*, *M. nigra*, and *M. pleuralis*) fungus-feeding larvae are overwhelmingly common. Sporophore eruptions can produce hundreds of flies relatively quickly, as Brown & Hartop [58,59] estimated a single mushroom cap contained 500 larvae of *M. marquezi.* We suspect abundance peaks for these species are associated with the mass production of fungal sporophores in close proximity to our sampling area, which are common after rain. However, we also suspect that very wet climates may increase fungal disease incidence [60,61], such that highest phorid fly richness occurs at intermediate relative humidity.

Urban landscapes are rarely designed to sustain biodiversity, though this is often an idealized goal informed by research [62]. At very high levels of heat, abundance and diversity declines, which is consistent with other studies suggesting that the urban heat island effect has negative effects on many species [14,63,64]. As the global temperature increases, many of the sites we sampled may also warm and therefore no longer support diverse phorid fly communities, though this will depend on how quickly phorid flies can adapt to changing thermal conditions. Even on short timescales, it is possible that the thermal limits of species have evolved, so that animals in the warmer parts of the city are capable of tolerating warmer conditions independent of ecotypic acclimation [65]. Given the rapid evolution of thermal tolerance in other arthropods [65–67], and the short generation times of phorid flies, rapid adaptation to climate change might be possible.

Our work was designed to test how the urban matrix surrounding sites in urban Los Angeles affects insect biodiversity. Our analyses suggest that temperature is a more important variable than surrounding urban land cover (impervious surface and NDVI densities) for determining fly abundance and richness. However, a more detailed analysis of the specific habitat types between potential source communities in protected, more natural sites and urban sites might reveal patterns we have not tested for. Specifically, we predict that if protected areas are source populations for urban fly biodiversity, protected areas and urban sites with more hospitable habitat between (more NDVI, for example), may have higher fly diversity. Assessing the specific land cover types between protected areas and urban sites is an important area for future studies. We also note that our study did not take into account plant species composition, including the amount of native vs. non-native plant cover. Prior work has shown that the amount of native vegetation [68], vegetation complexity [69,70], and plant diversity [71] can drive urban insect diversity. Investigating plant species composition of the matrix around sites and intra-urban corridors among sites may help explain patterns in diversity that are not accounted for in our analyses.Exhaustive biodiversity sampling has reaped substantial rewards in understanding how environmental change across space affects biodiversity [72,73]. While labor intensive, our approach created a foundational understanding of which species occur in the phorid fly community, a presumably informative subset of the entire insect community. Baseline knowledge of insect communities is a prerequisite for generating expected responses to continued global change. These data are rare, but sampling programs like the one we describe here could be replicated in other cities to build baselines that allow us to determine how biotic change varies across background climates and habitat types. However, our robust sampling of the L.A. Basin relied on collaboration between scientists and the public. Members of the community hosted Malaise traps in their backyards, increasing the range of urban environments available for sampling, and, importantly, reducing the resources and labor required for this intensive sampling [74]. With continued public support, efforts such as ours could create long-term data for describing species’ long-term responses to urbanization and climate change [75,76].

## Supporting information

Electronic Supplementary Material, including figures and some methods

## Authors’ Contributions

The BioSCAN project was conceived and operated by BVB, and taxonomy and identification of flies was conducted by BVB and EAH. Data curation and analyses were conducted by EKM and CAB. Spatial data for analyses and maps were provided by EL. The manuscript was written by TPM, with contributions from EKM, BVB, BJA, EL, and CAB.

## Acknowledgments

BioSCAN is a large project, funded by the LACM, its donors, and grants from the Seaver Foundation and the Rose Hills Foundation. We thank the many staff, supporters, and volunteers who have made the project possible, especially Luis Chiappe, Victoria Dean, Esther Chao, Lisa Gonzalez, Estella Hernandez, Lila Higgins, Tom Jacobson, Diane Naegele, Greg Pauly, Dean Pentcheff, Chris Thacker, Jann Vendetti, Regina Wetzer, and Maria Wong. We thank the families and representatives of organizations whose homes or yards were made available to us for sampling, including Celeste Armstrong, Natalie Brejcha, Chris Cianci, Glen Creason, Teresa Dahl, Betty Defibaugh, Julian Donahue, Candace Franco, Ray Fujioka, Victoria Harding, Tony Hein, Peggy Hentschke, Sidney Higgins, Jim Hogue, Jeff Koch, Erin Johnson, Eric Keller, Patricia Lombard, Pauline Louie, Humberto Marquez, Sharon Oxborough, Mark Pisano, Peter Ralph, Walter Renwick, John Rodriguez, LaChristian Steptoe, and K.T. Wiegman. TPM was supported by a sabbatical and RSCA awards from CSUDH and by NSF (OISE-1261015)

## Notes

#### Summary of Updates

This is a revised version of the manuscript after peer review, with new supplemental figures added and several clarifications have been added to the text.

https://doi.org/10.5061/dryad.gr68f2j

## References

1. Lundholm JT, Marlin A. 2006 Habitat origins and microhabitat preferences of urban plant species. Urban Ecosyst. (doi:10.1007/s11252-006-8587-4)

2. Aronson MFJ et al. 2014 A global analysis of the impacts of urbanization on bird and plant diversity reveals key anthropogenic drivers. Proc. R. Soc. B Biol. Sci. (doi:10.1098/rspb.2013.3330)

3. McKinney ML. 2002 Urbanization, Biodiversity, and Conservation. Bioscience (doi:10.1641/0006-3568(2002)052[0883:UBAC]2.0.CO;2)

4. Gómez-Baggethun E et al. 2013 Urban Ecosystem Services. In Urbanization, Biodiversity and Ecosystem Services: Challenges and Opportunities, (doi:10.1007/978-94-007-7088-1)

5. Potter A, LeBuhn G. 2015 Pollination service to urban agriculture in San Francisco, CA. Urban Ecosyst. (doi:10.1007/s11252-015-0435-y)

6. Alberti M. 2005 The effects of urban patterns on ecosystem function. Int. Reg. Sci. Rev.(doi:10.1177/0160017605275160)

7. Shochat E, Warren PS, Faeth SH, McIntyre NE, Hope D. 2006 From patterns to emerging processes in mechanistic urban ecology. Trends Ecol. Evol. (doi:10.1016/j.tree.2005.11.019)

8. Lin BB, Philpott SM, Jha S. 2015 The future of urban agriculture and biodiversity-ecosystem services: Challenges and next steps. Basic Appl. Ecol. (doi:10.1016/j.baae.2015.01.005)

9. Yuan F, Bauer ME. 2007 Comparison of impervious surface area and normalized difference vegetation index as indicators of surface urban heat island effects in Landsat imagery. Remote Sens. Environ. 106, 375–386. (doi:https://doi.org/10.1016/j.rse.2006.09.003)

10. Meineke EK, Dunn RR, Frank SD. 2014 Early pest development and loss of biological control are associated with urban warming. Biol. Lett. 10. (doi:10.1098/rsbl.2014.0586)

11. Meineke EK, Frank SD. 2018 Water availability drives urban tree growth responses to herbivory and warming. J. Appl. Ecol. (doi:10.1111/1365-2664.13130)

12. Meineke E, Youngsteadt E, Dunn RR, Frank SD. 2016 Urban warming reduces aboveground carbon storage. Proc. R. Soc. B Biol. Sci. (doi:10.1098/rspb.2016.1574)

13. Meineke EK, Holmquist AJ, Wimp GM, Frank SD. 2017 Changes in spider community composition are associated with urban temperature, not herbivore abundance. J. Urban Ecol. (doi:10.1093/jue/juw010)

14. Hamblin AL, Youngsteadt E, López-Uribe MM, Frank SD. 2017 Physiological thermal limits predict differential responses of bees to urban heat-island effects. Biol. Lett. 13.

15. Dale AG, Frank SD. 2018 Urban plants and climate drive unique arthropod interactions with unpredictable consequences. Curr. Opin. Insect Sci. (doi:https://doi.org/10.1016/j.cois.2018.06.001)

16. Oke TR. 1973 City size and the urban heat island. Atmos. Environ. 7, 769–779. (doi:https://doi.org/10.1016/0004-6981(73)90140-6)

17. IPCC. 2014 Summary for Policymakers. (doi:10.1017/CBO9781107415324)

18. Meineke EK, Dunn RR, Sexton JO, Frank SD. 2013 Urban Warming Drives Insect Pest Abundance on Street Trees. PLoS One 8, 1–7. (doi:10.1371/journal.pone.0059687)

19. Dale AG, Frank SD. 2014 Urban warming trumps natural enemy regulation of herbivorous pests. Ecol. Appl. (doi:10.1890/13-1961.1)

20. Tibbetts EA, Dale J. 2004 A socially enforced signal of quality in a paper wasp. 432, 218–222. (doi:10.1038/nature03004.1.)

21. Dale AG, Frank SD. 2017 Warming and drought combine to increase pest insect fitness on urban trees. PLoS One (doi:10.1371/journal.pone.0173844)

22. Neil K, Wu J. 2006 Effects of urbanization on plant flowering phenology: A review. Urban Ecosyst. (doi:10.1007/s11252-006-9354-2)

23. Somers KA, Bernhardt ES, Grace JB, Hassett BA, Sudduth EB, Wang S, Urban DL. 2013 Streams in the urban heat island: spatial and temporal variability in temperature. Freshw. Sci. (doi:10.1899/12-046.1)

24. Faeth SH, Bang C, Saari S. 2011 Urban biodiversity: Patterns and mechanisms. Ann. N. Y. Acad. Sci. (doi:10.1111/j.1749-6632.2010.05925.x)

25. Taha H. 1997 Urban climates and heat islands: albedo, evapotranspiration, and anthropogenic heat. Energy Build. (doi:10.1016/S0378-7788(96)00999-1)

26. Luo Z, Sun OJ, Ge Q, Xu W, Zheng J. 2007 Phenological responses of plants to climate change in an urban environment. Ecol. Res. (doi:10.1007/s11284-006-0044-6)

27. Youngsteadt E, Dale AG, Terando AJ, Dunn RR, Frank SD. 2015 Do cities simulate climate change? A comparison of herbivore response to urban and global warming. Glob. Chang. Biol. 21, 97–105. (doi:10.1111/gcb.12692)

28. Lahr EC, Dunn RR, Frank SD. 2018 Getting ahead of the curve: cities as surrogates for global change. In Proceedings of the Royal Society B: Biological Sciences, (doi:10.1098/rspb.2018.0643)

29. Melillo JM et al. 2011 Soil warming, carbon-nitrogen interactions, and forest carbon budgets. Proc. Natl. Acad. Sci. (doi:10.1073/pnas.1018189108)

30. Rinnan R, Michelsen A, Bååth E, Jonasson S. 2007 Fifteen years of climate change manipulations alter soil microbial communities in a subarctic heath ecosystem. Glob. Chang. Biol. (doi:10.1111/j.1365-2486.2006.01263.x)

31. Weng Q, Lu D, Schubring J. 2004 Estimation of land surface temperature-vegetation abundance relationship for urban heat island studies. Remote Sens. Environ. (doi:10.1016/j.rse.2003.11.005)

32. Cadenasso ML, Pickett STA, Schwarz K. 2007 Spatial heterogeneity in urban ecosystems: Reconceptualizing land cover and a framework for classification. Front. Ecol. Environ. (doi:10.1890/1540-9295(2007)5[80:SHIUER]2.0.CO;2)

33. Buyantuyev A, Wu J. 2010 Urban heat islands and landscape heterogeneity: Linking spatiotemporal variations in surface temperatures to land-cover and socioeconomic patterns. Landsc. Ecol. (doi:10.1007/s10980-009-9402-4)

34. McIntyre NE. 2000 Ecology of Urban Arthropods: A Review and a Call to Action. Ann. Entomol. Soc. Am. (doi:10.1603/0013-8746(2000)093[0825:EOUAAR]2.0.CO;2)

35. Gardiner MM, Prajzner SP, Burkman CE, Albro S, Grewal PS. 2014 Vacant land conversion to community gardens: Influences on generalist arthropod predators and biocontrol services in urban greenspaces. Urban Ecosyst. (doi:10.1007/s11252-013-0303-6)

36. Borkent A et al. 2018 Remarkable fly (Diptera) diversity in a patch of Costa Rican cloud forest: Why inventory is a vital science. Zootaxa. (doi:10.11646/zootaxa.4402.1.3)

37. Brown B V. et al. 2018 Comprehensive inventory of true flies (Diptera) at a tropical site. Commun. Biol. (doi:10.1038/s42003-018-0022-x)

38. Disney RHL. 1990 Some myths and the reality of scuttle fly biology. Antenna

39. Hartop EA, Brown B V, Disney RHL. 2015 Opportunity in our Ignorance: Urban Biodiversity Study Reveals 30 New Species and One New Nearctic Record for Megaselia (Diptera: Phoridae) in Los Angeles (California, USA). Zootaxa; Vol 3941, No 4 2 Apr. 2015 (doi:10.11646/zootaxa.3941.4.1)

40. Hartop EA, Brown B V, Disney RHL. 2016 Flies from L.A., The Sequel: A further twelve new species of Megaselia (Diptera: Phoridae) from the BioSCAN Project in Los Angeles (California, USA). Biodivers. Data J., e7756. (doi:10.3897/BDJ.4.e7756)

41. Vahmani P, Ban-Weiss GA. 2016 Impact of remotely sensed albedo and vegetation fraction on simulation of urban climate in WRF-urban canopy model: A case study of the urban heat island in Los Angeles. J. Geophys. Res. (doi:10.1002/2015JD023718)

42. Brown B V, Hartop EA. 2017 Big data from tiny flies: patterns revealed from over 42,000 phorid flies (Insecta: Diptera: Phoridae) collected over one year in Los Angeles, California, USA. Urban Ecosyst. 20, 521–534. (doi:10.1007/s11252-016-0612-7)

43. Melles S, Glenn S, Martin K. 2003 Urban bird diversity and landscape complexity: species–environment associations along a multiscale habitat gradient. Conserv. Ecol. 7.

44. Keeley JE, Swift CC. 2011 Biodiversity and Ecosystem Functioning in Mediterranean-Climate California. (doi:10.1007/978-3-642-78881-9_3)

45. Roth M, Oke TR, Emery WJ. 1989 Satellite-derived urban heat islands from three coastal cities and the utilization of such data in urban climatology. Int. J. Remote Sens. (doi:10.1080/01431168908904002)

46. Witiw MR, LaDochy S. 2008 Trends in fog frequencies in the Los Angeles Basin. Atmos. Res. (doi:10.1016/j.atmosres.2007.11.010)

47. Townes H. 1972 A light-weight Malaise trap. Entomol. News 83, 239–247.

48. Cayelan CC, Cottingham KL. 2016 Cross-scale Perspectives: Integrating Long-term and High-frequency Data into Our Understanding of Communities and Ecosystems. Bull. Ecol. Soc. Am. 97, 129–132. (doi:10.1002/bes2.1205)

49. NOAA. 2019 Climate Data Online. See https://www.ncdc.noaa.gov/cdo-web/ (accessed on 20 September 2001).

50. Warren II RJ, Bayba S, Krupp KT. 2018 Interacting effects of urbanization and coastal gradients on ant thermal responses. J. Urban Ecol. (doi:10.1093/jue/juy026)

51. Hartop EA, Gonzalez LA, Brown B V. 2018 Backyard biodiversity: Unraveling life histories of the new fly species discovered by the BioSCAN Project proves harder than first assumed. J. Negat. Results 12.

52. Pinheiro J, Bates D, DebRoy S, Sarkar D. 2007 nlme: Linear and Nonlinear Mixed Effects Models.

53. Trumble JT, Pienkowski RL. 1979 Development and survival of Megaselia scalaris (Diptera: Phoridae) at selected temperatures and photoperiods. Proc Entomol Soc Wash 81, 207–210.

54. Paradis E, Claude J, Strimmer K. 2004 APE: Analyses of phylogenetics and evolution in R language. Bioinformatics (doi:10.1093/bioinformatics/btg412)

55. Oksanen J et al. 2013 Vegan: community ecology package. R Packag. Version 2.9 (doi:10.1093/molbev/msv334)

56. Poos MS, Jackson DA. 2012 Addressing the removal of rare species in multivariate bioassessments: The impact of methodological choices. Ecol. Indic. (doi:10.1016/j.ecolind.2011.10.008)

57. Suarez A V, Case TJ. 2003 9 The Ecological Consequences of a Fragmentation Mediated Invasion: The Argentine Ant, Linepithema humile, in Southern California. 162.

58. Hartop E, Brown B. 2014 The tip of the iceberg: a distinctive new spotted-wing Megaselia species (Diptera: Phoridae) from a tropical cloud forest survey and a new, streamlined method for Megaselia descriptions. Biodivers. Data J. (doi:10.3897/BDJ.2.e4093)

59. Brown B V, Hartop EA. 2017 Mystery mushroom malingerers: Megaselia marquezi Hartop et al. 2015 (Diptera: Phoridae). Biodivers. Data J. (doi:10.3897/BDJ.5.e15052)

60. Araújo JPM, Hughes DP. 2016 Diversity of Entomopathogenic Fungi. Which Groups Conquered the Insect Body? In Advances in Genetics, (doi:10.1016/bs.adgen.2016.01.001)

61. Ramoska WA. 1984 The influence of relative humidity on Beauveria bassiana infectivity and replication in the chinch bug, Blissus leucopterus. J. Invertebr. Pathol. (doi:10.1016/0022-2011(84)90085-5)

62. Dunn RR, Gavin MC, Sanchez MC, Solomon JN. 2006 The pigeon paradox: Dependence of global conservation on urban nature. Conserv. Biol. (doi:10.1111/j.1523-1739.2006.00533.x)

63. Dale AG, Frank SD. 2018 Urban plants and climate drive unique arthropod interactions with unpredictable consequences. Curr. Opin. Insect Sci. (doi:10.1016/J.COIS.2018.06.001)

64. Youngsteadt E, Ernst AF, Dunn RR, Frank SD. 2017 Responses of arthropod populations to warming depend on latitude: evidence from urban heat islands. Glob. Chang. Biol. (doi:10.1111/gcb.13550)

65. Diamond SE, Chick L, Perez A, Strickler SA, Martin RA. 2017 Rapid evolution of ant thermal tolerance across an urban-rural temperature cline. Biol. J. Linn. Soc. 121, 248–257. (doi:10.1093/biolinnean/blw047)

66. I. BK, Luc DM. 2018 City life on fast lanes: urbanization induces an evolutionary shift towards a faster life style in the water flea Daphnia. Funct. Ecol. 0. (doi:10.1111/1365-2435.13184)

67. Tüzün N, Op de Beeck L, Brans KI, Janssens L, Stoks R. 2017 Microgeographic differentiation in thermal performance curves between rural and urban populations of an aquatic insect. Evol. Appl. (doi:10.1111/eva.12512)

68. Burghardt KT, Tallamy DW, Gregory Shriver W. 2009 Impact of native plants on bird and butterfly biodiversity in suburban landscapes. Conserv. Biol. (doi:10.1111/j.1523-1739.2008.01076.x)

69. Beninde J, Veith M, Hochkirch A. 2015 Biodiversity in cities needs space: A meta- analysis of factors determining intra-urban biodiversity variation. Ecol. Lett. (doi:10.1111/ele.12427)

70. Snep RPH, Opdam PFM, Baveco JM, WallisDeVries MF, Timmermans W, Kwak RGM, Kuypers V. 2006 How peri-urban areas can strengthen animal populations within cities: A modeling approach. Biol. Conserv. (doi:10.1016/j.biocon.2005.06.034)

71. Kearns CA, Oliveras DM. 2009 Environmental factors affecting bee diversity in urban and remote grassland plots in Boulder, Colorado. J. Insect Conserv. (doi:10.1007/s10841-009-9215-4)

72. Longino JT, Colwell RK. 1997 Biodiversity assessment using structured inventory: Capturing the ant fauna of a tropical rain forest. Ecol. Appl. (doi:10.1890/1051-0761(1997)007[1263:BAUSIC]2.0.CO;2)

73. Hallmann CA et al. 2017 More than 75 percent decline over 27 years in total flying insect biomass in protected areas. PLoS One (doi:10.1371/journal.pone.0185809)

74. Dickinson JL, Zuckerberg B, Bonter DN. 2010 Citizen Science as an Ecological Research Tool: Challenges and Benefits. Annu. Rev. Ecol. Evol. Syst. (doi:10.1146/annurev-ecolsys-102209-144636)

75. Wolkovich EM, Regetz J, O’Connor MI. 2012 Advances in global change research require open science by individual researchers. Glob. Chang. Biol. (doi:10.1111/j.1365-2486.2012.02693.x)

76. Theobald EJ et al. 2015 Global change and local solutions: Tapping the unrealized potential of citizen science for biodiversity research. Biol. Conserv. (doi:10.1016/j.biocon.2014.10.021)

