## Supplementary material for "Temperature accounts for the biodiversity of a hyperdiverse group of insects in urban Los Angeles": Electronic Supplementary Material, including figures and some methods

*Methods: Abiotic data collection and processing*

Weather stations were regularly maintained, but like most sensor-based ecological monitoring, gaps in reading due to occasional equipment failure were common [48]. We addressed these gaps through a series of down-sampling and estimation methods. First, data for readings of each type were subjected to quality control to remove impossible values, and flag improbable values for additional scrutiny. Then, to eliminate small gaps in the data, because readings were taken at five-minute intervals, we averaged all readings for each measure by hour. We culled out observations from time periods where fewer than 18 of the 30 sites were reporting data (primarily occurring during experimental set-up and take down). Following this process, null observations (gaps) represented an average of 4% of site-by-hour observations, and 11 of the 30 sites had no gaps. The weather in the year of this study (2014) was hotter than previous years, but not particularly anomalous relative to recent years [49]. Because many sites were near to each other and behaved similarly (although not identically) from an environmental standpoint, we estimated missing data first by using principal component analysis to rank sites, pairwise, by their similarity. Then, for each site with missing values for a given environmental response variable, we ran a linear model between that variable and the same variable from the most similar site. Then, based on the regression parameters observed, we estimated the values of the missing data, and substituted them for the missing value code into the weather data frame. We repeated this process for each site and variable combination where missing values were observed, and in the rare occasion where this process did not address all the gaps, we repeated this process by pairing the site with missing data from the second most similar site.

We estimated distance to the closest natural area by calculating the distance to the nearest boundary of an area recorded in the California Protected Area Database (CPAD). CPAD contains lands protected under conservation easements and open space, such as national/state/regional parks, forests, preserves, and wildlife areas. As estimates of urbanization, we used 2016 -2017 U.S. Geological Survey Landsat 7 surface reflectance and the National Land Cover Database 2011 Percent Developed Imperviousness Layer to calculate the average Normalized Difference Vegetation Index (NDVI) and the percentage of impervious surface at each site. A 50-meter buffer was used for data extraction to represent the local environment.

Most non-climatic variables included in this study remained fairly constant over the study period. These variables could be represented as single values per site in our models (distance to CPAD, percent impervious surface). However, NDVI may shift throughout the year. To assess the possibility that variation in NDVI was substantial enough to require us to represent changing NDVI throughout the year in our models, we used a google API tool (https://clim-engine.appspot.com) to generate monthly NDVI means for each month during 2014—the year of our sampling—based on Landsat 8 imagery, which is at 30-meter resolution. We plotted monthly NDVI means for each site in each month (Fig. S2). We also plotted mean NDVI with standard errors representing variation within the year 2014 across our study sites (Fig. S3). We found that variation within the year across sites was minimal and therefore used summary NDVI values as described above in analyses.

Figure S1. Scatterplot matrix showing correlations between environmental and microclimatic predictors: Mean weekly air temperature (ºC), average normalized difference vegetation index for a circle with radius of 50 m surrounding each site (NDVI), percent impervious groundcover in that same area and distance to nearest natural area (m). Straight pale green lines represent the robust-regression line between the variable pairs, and the curved red lines represent a GAM (generalized additive model) smoother and their respective confidence intervals.

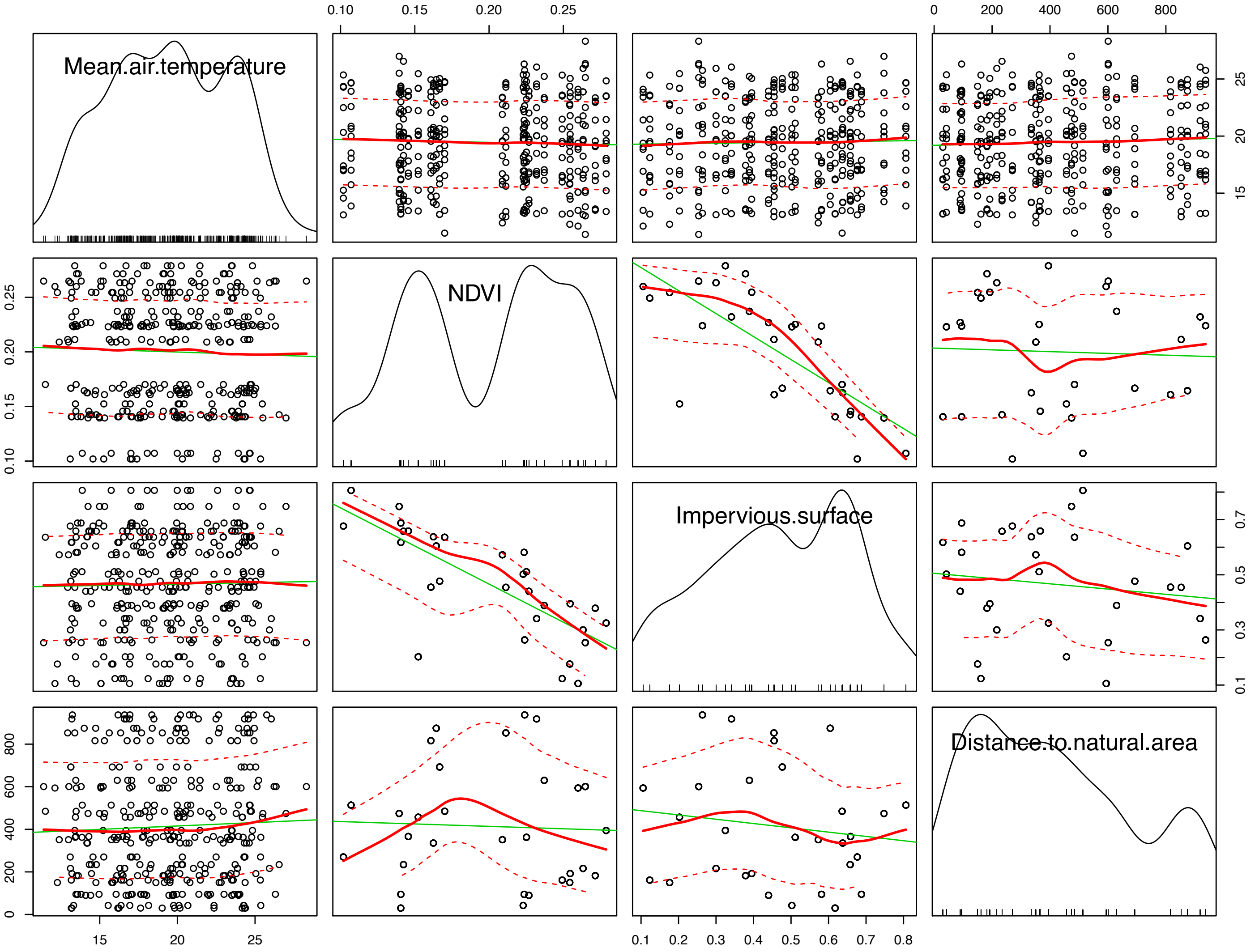

Figure S2. Mean Normalized Difference Vegetation Index (NDVI), representing vegetation cover surrounding each site, along with standard error bars.

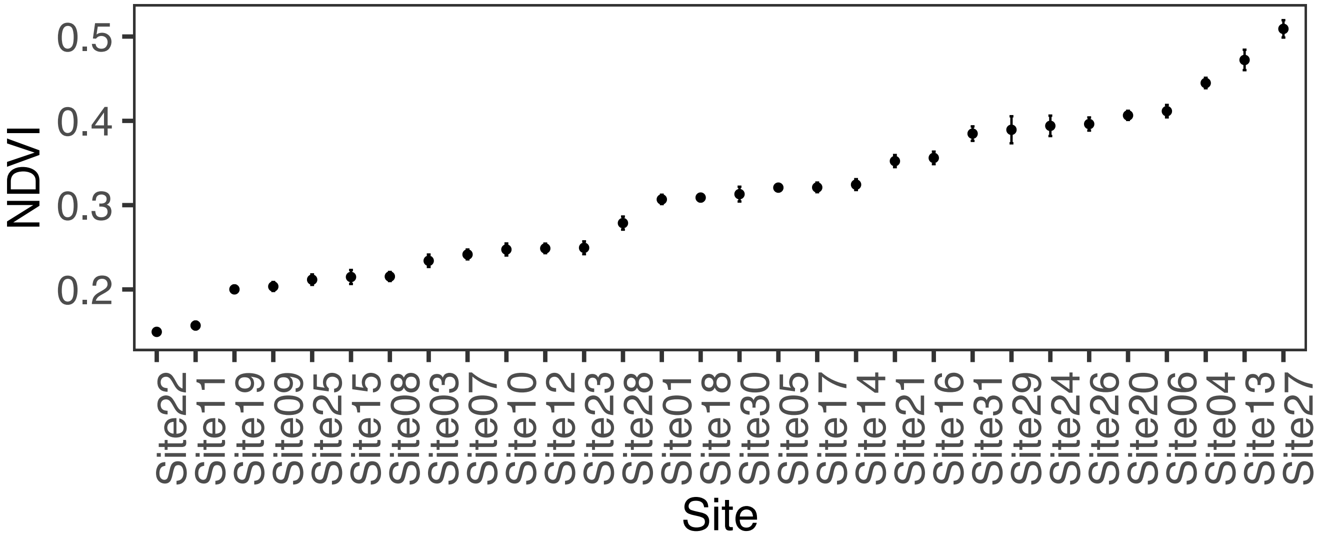

Figure S3. Scatterplot matrix of relative humidity, minimum soil temperature (ºC) at 1-cm depth, mean weekly air temperature (ºC), and distance to nearest natural, protected area (m). Straight pale green lines represent the robust-regression line between the variable pairs, and the curved red lines represent a GAM smoother and their respective confidence intervals.

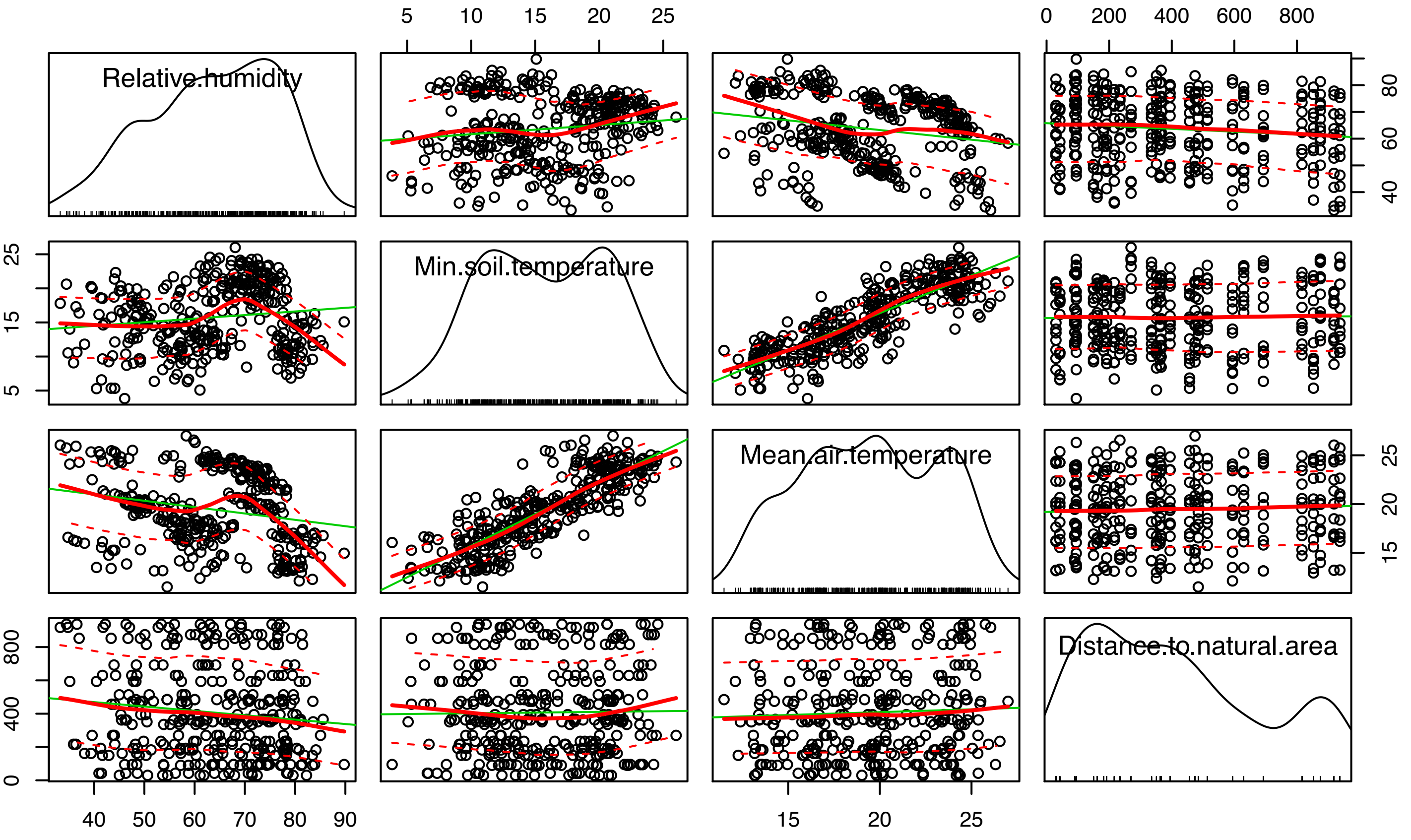

Figure S4. Scatterplot matrix of latitude, longitude, relative humidity, minimum soil temperature (ºC) at 1-cm depth, and mean air temperature (ºC). Straight pale green lines represent the robust-regression line between the variable pairs, and the curved red lines represent a GAM smoother and their respective confidence intervals.

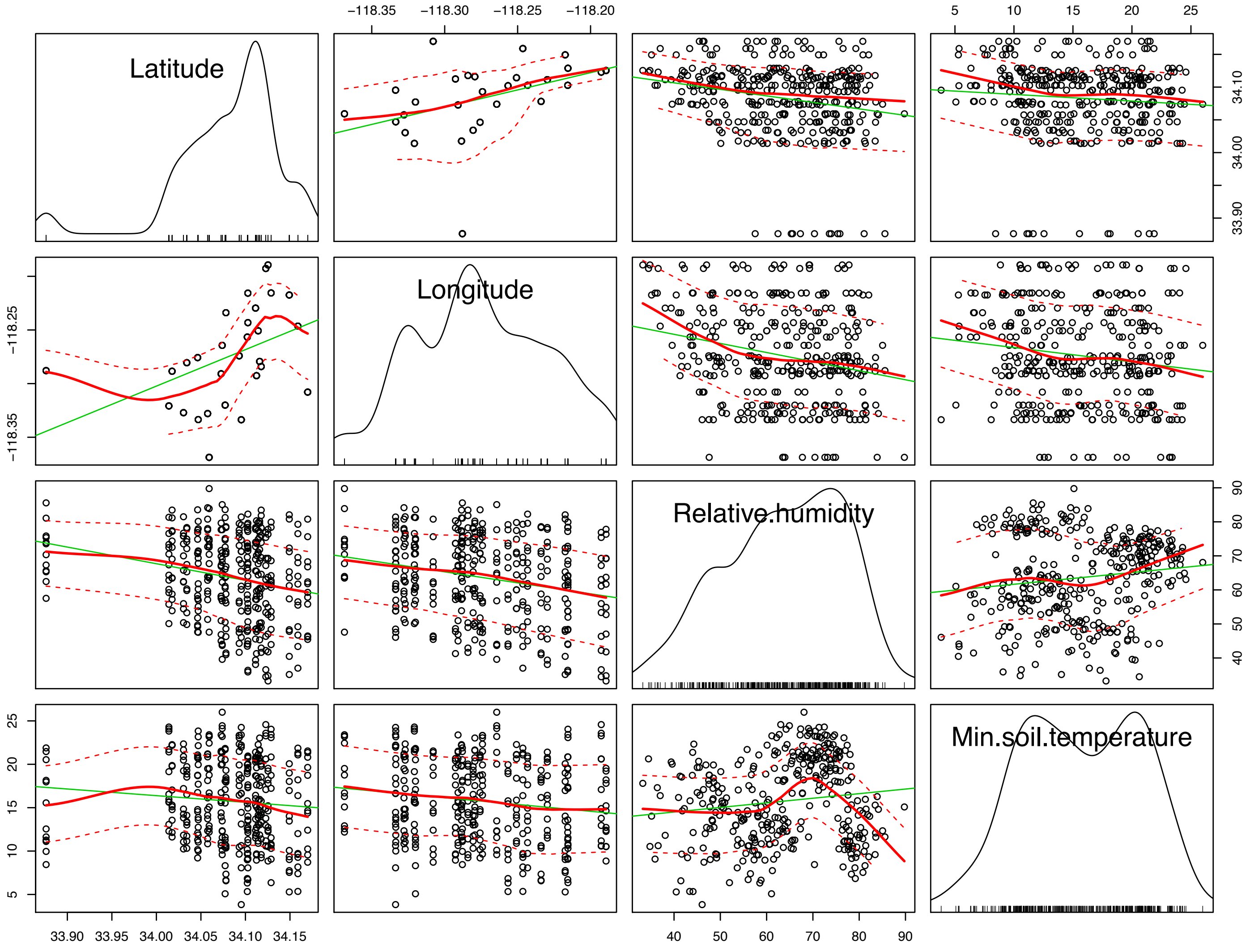

Figure S5. Site maps of abiotic variables analyzed, including a) NDVI (50-meter buffer), b) impervious surface (50-m buffer), and c) distance to closest natural, protected area.

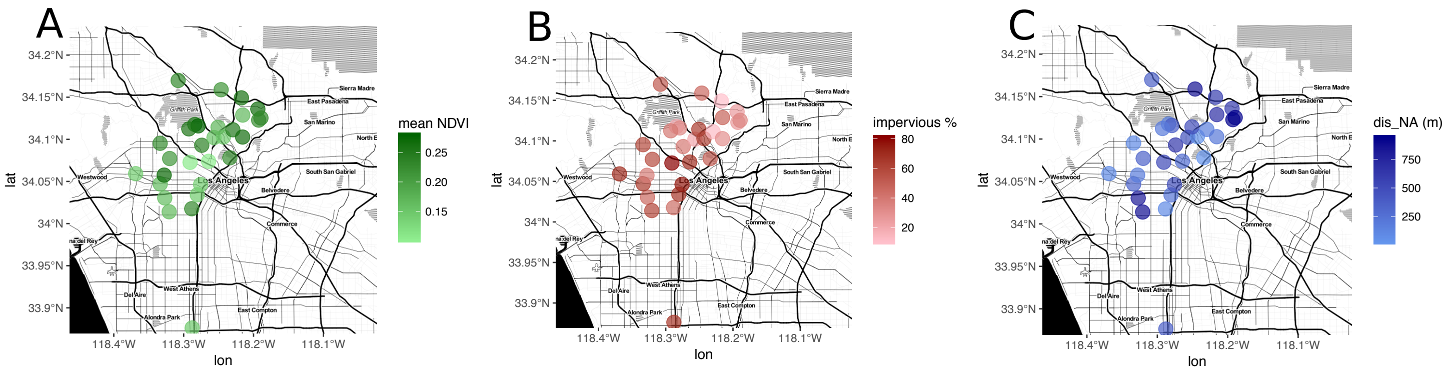

Figure S6. Variation in the mean Normalized Difference Vegetation Index (NDVI), representing vegetation cover surrounding each site, over 2014, the year phorid flies were sampled.

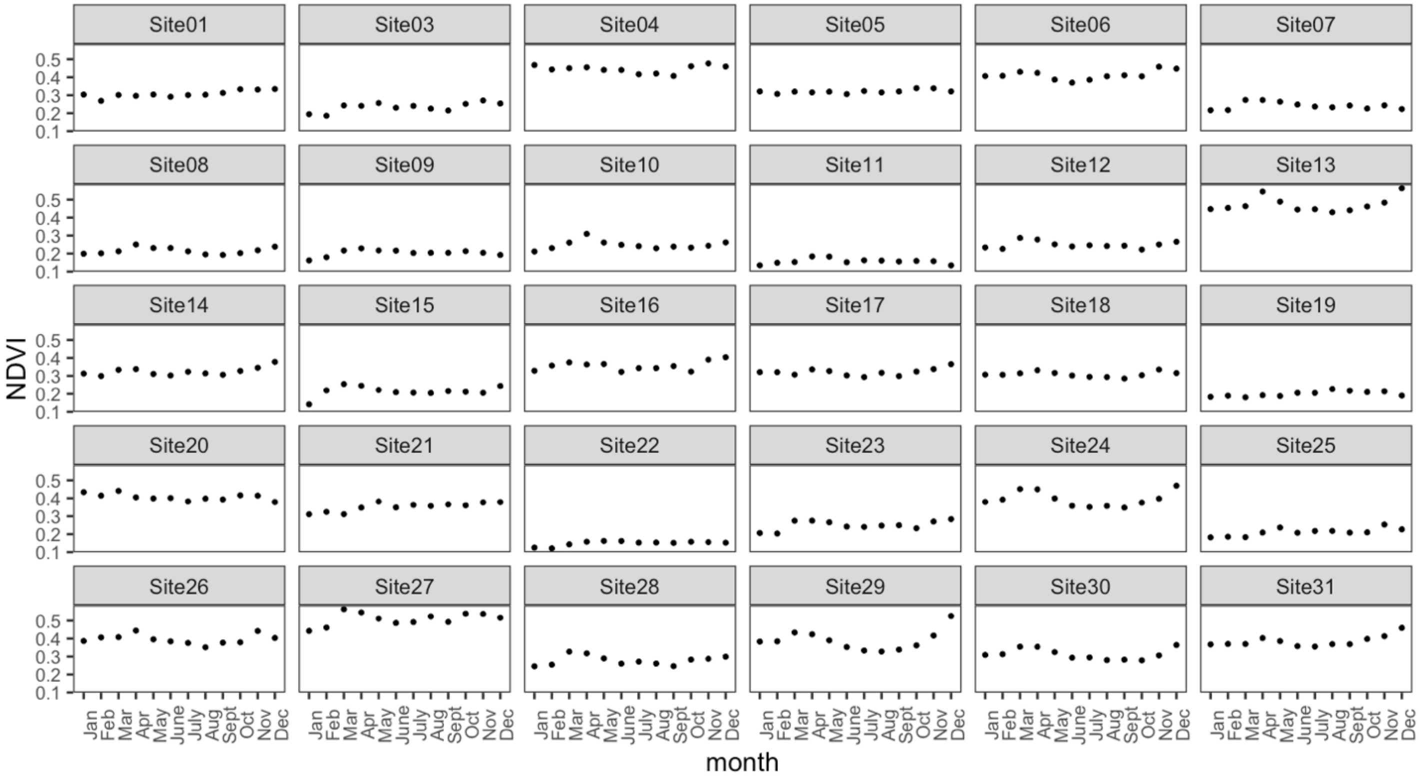
